# CRISPR-based bioengineering of the Transferrin Receptor revealed a role for Rab7 in the biosynthetic secretory pathway

**DOI:** 10.1101/2020.01.05.893206

**Authors:** Maika S. Deffieu, Ieva Cesonyte, François Delalande, Gaelle Boncompain, Cristina Dorobantu, Eli Song, Vincent Lucansky, Aurélie Hirschler, Sarah Cianferani, Tao Xu, Franck Perez, Christine Carapito, Raphael Gaudin

## Abstract

The regulated secretory trafficking of neosynthesized transmembrane receptors is particularly challenging to investigate as it is under-represented at steady state compared to the abundance of the other trafficking routes. Here, we combined the retention using selective hook (RUSH) system to a CRISPR/Cas9 gene editing approach (eRUSH) to identify molecular players involved in the trafficking of neosynthesized Transferrin Receptor (TfR) *en route* to the plasma membrane (PM). TfR-eRUSH monoclonal cells expressing endogenous, ER-retainable and fluorescent TfR were engineered and characterized. Spatiotemporal quantitative proteomics of TfR-eRUSH cells allowed the identification of molecular partners associated with TfR-containing membranes and provided a comprehensive list of potential regulators, co-trafficking cargos, and enriched pathways. Furthermore, we chose to focus our attention on the Rab GTPase family members for their function as vesicle trafficking regulators and performed a Rab-targeted siRNA screen that we correlated to our proteomics data. Unexpectedly, we identified Rab7-harboring vesicles as an intermediate compartment of the Golgi-to-PM transport of the neosynthetic TfR. These vesicles did not exhibit degradative properties and were not associated to Rab6A-harboring vesicles, also involved in Golgi-to-PM transport. However, Rab6A-TfR vesicles delivered TfR directly to the PM, while in contrast, Rab7A was transiently associated to neosynthetic TfR-containing post-Golgi vesicles but dissociated before PM vesicle fusion. Together, our study proposes the eRUSH as a powerful tool to further study the secretory pathway and reveals an unforeseen role for Rab7 in the neosynthetic transport of the TfR, highlighting the diversity of the secretory vesicles’ nature for a given cargo.

## Introduction

Cells sense environmental changes and adapt accordingly by exposing a variety of transmembrane receptors at their cell surface. Post-translational modifications and final localization of these transmembrane receptors at the plasma membrane (PM) are first occurring through the membrane dynamics along the secretory pathway. The secretory pathway is a constitutive or regulated process (1) carrying neosynthesized proteins from the endoplasmic reticulum (ER) to the PM. Characterizing the molecular mechanisms involved in this cellular process may be useful for the development of inhibitors targeting general or cargo-specific secretion (2).

Transmembrane receptors are synthesized and folded in the ER. After synthesis, coatomer protein complex II (COP-II) vesicles export the incorporated receptors to the *cis*-Golgi cisternae (3). The transit of these cargoes through the Golgi stacks is still debated (4, 5), although it is well established that proteins undergo successive post-translational modifications during their trafficking from the *cis*-Golgi to the TGN. Upon protein arrival at the TGN, cargoes are specifically packaged and sorted to be delivered to different organelles such as endosomes, lysosomes or the PM. Sorting signals identified at the cytosolic regions of transmembrane receptors lead to the specific recruitment of adaptor proteins (APs) or small Rab-GTPases, needed for the incorporation of the cargo inside vesicle carriers. After budding off the TGN membranes, proteins are delivered to their final destination through vesicular transport. It was long being thought that transmembrane receptors use a direct route from the TGN to the PM. Observations of differential trafficking routes suggested otherwise. Indeed, several studies noticed the presence of cargoes inside endocytic compartments before their delivery to the PM (6–8). The nature and fate of these intermediate compartments in protein secretion is still unclear.

To mechanistically understand temporally and spatially the secretory pathway, a few systems were developed. One of the earliest methods developed to study protein secretion was the thermo-sensitive vesicular stomatitis virus glycoprotein (ts045VSV-G) (9). It involves incubation of cells at a restrictive temperature to block ts045VSV-G transport at the ER followed by a shift at a lower permissive temperature to induce the release of the protein to its normal trafficking pathway (10). This method provided valuable analytical information on the dynamics and kinetics of transport of ts045VSV-G from the TGN to the PM.

To avoid non-physiological conditions of temperature and monitor different cargo proteins, the RUSH (retention using selective hooks) system was elaborated (11). It allows the retention of a protein of interest in the ER, then its on-demand release following the addition of biotin in the cell media. This method proved to be very powerful (2,12–15), but it requires the transient overexpression of the protein of interest which is a limitation in case of regulated secretion. In addition, the co-existence of the overexpressed tagged and the non-tagged endogenous cargos could confer some limitations for quantitative temporal detection of a receptor at the PM.

The vesicular carriers involved in the secretory pathway are difficult to study because of their low abundance at steady state compared to endocytic/recycling vesicles. This is particularly true for the Transferrin Receptor 1 (TfR), which is widely used for recycling studies (for review see (16)). TfR is a ubiquitous transmembrane glycoprotein that mediates iron uptake from circulating transferrin (Tf) at the PM. After formation of the TfR-Tf complex at the cell surface, the receptor is internalized by clathrin-mediated endocytosis and delivered to endosomes. Inside these organelles, TfR dissociates from its ligand and is recycled back to the cell surface. Studies indicated that an alteration of the expression level of TfR could trigger carcinoma progression (17, 18). Indeed, cancer cells expressed a high amount of TfR at their cell surface which makes it a significant anti-cancer target (19, 20).

Neosynthesized TfR arriving at the PM represents a minor fraction of the total TfR pool expressed at the cell surface at steady state, and thus the pathway of newly synthesized TfR is particularly difficult to investigate. In this study, we developed an approach that combines the RUSH system with CRISPR/Cas9 gene editing that we called “edited-RUSH” or “eRUSH”. We employed eRUSH to investigate the molecular mechanisms involved in the vesicular transport of neosynthesized TfR to the PM. The TfR-eRUSH allowed the spatiotemporal monitoring of the trafficking of the neosynthesized endogenous TfR as well as the identification of the molecular partners involved in this process. In particular, we highlighted that Rab7A, a small Rab GTPase usually described as an endolysosomal marker, is required for efficient arrival of neosynthesized TfR at the PM and was recruited to a subset of post-TGN TfR-containing vesicles, suggesting that Rab7 may play a role in the anterograde trafficking pathway of secretory vesicles.

## Results

### Generation and characterization of the TfR-eRUSH system

The CRISPR/Cas9 strategy that we previously described (21) was used to engineer the breast cancer-derived SUM159 cells to express endogenous TfR fused to the streptavidin-binding peptide (SBP) and EGFP. Lentiviral transduction of a chimera streptavidin-KDEL protein was performed to establish a stable cell line that retains SBP-containing proteins in the ER (see (11) for original description of the RUSH system) and the resulting TfR-eRUSH cells were subsequently characterized.

As depicted in Figure 1A, SBP fused to EGFP was introduced in the genomic sequence of TfR before its stop codon sequence. At the genomic DNA level, both alleles were carrying an extra piece of DNA corresponding to the SBP-EGFP tag (Figure 1B). At the protein level, almost no endogenous TfR was detected (at ≈ 84 kDa), while an upper band at ≈ 117 kDa appeared, corresponding to the expected size of TfR-SBP-EGFP protein (Figure S1A). Of note, the molecular weights were difficult to precisely assess as ladders from different brands were providing dissimilar sizes for a given band. Using an anti-EGFP antibody, we could confirm that TfR-SBP-EGFP was indeed running at an apparent size of 117 kDa (Figure S1B). Depending of the ladder used along our study, the TfR-eRUSH would appear as a band of either ≈ 98 kDa or 117 kDa, although both would correspond to the TfR-eRUSH.

**Fig 1.**
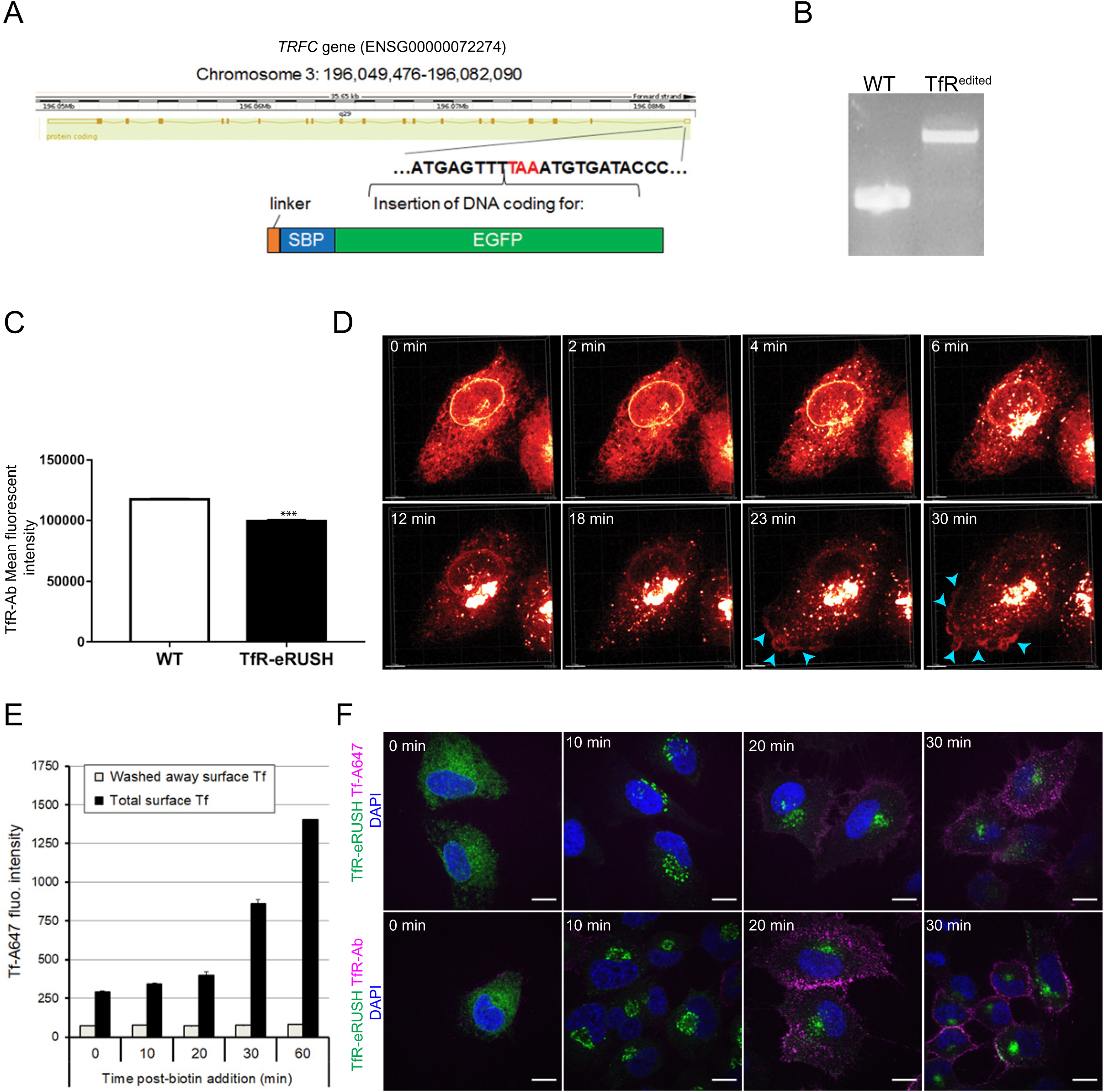
Generation and characterization of TfR-eRUSH gene edited cells. (A) Scheme illustrating the insertion of the linker-SBP-EGFP coding sequence in the chromosomal region containing the stop codon (red) of the *TFRC* gene (Transferrin receptor type 1, referred to as TfR). (B) PCR amplification from genomic DNA using primers flanking the TfR stop codon region confirmed the insertion of the SBP-EGFP sequence on both alleles. (C) Flow cytometry analysis indicates the total amount of TfR expressed in wild type (WT) and TfR-eRUSH cells, 24 h post-biotin treatment. Mouse anti-TfR antibody revealed with anti-mouse Alexa Fluor 647 antibody were used to measure total TfR protein levels. The bar graph indicates the mean fluorescence intensity +/- SD of cell populations expressing TfR. At least 10,000 cells per condition were acquired from n = 6 independent experiments. (D) Live cell imaging of TfR-eRUSH cells started immediately after biotin addition highlights the rapid and dramatic redistribution of TfR over time. Sequential trafficking steps include ER (0 to 6 min) to Golgi (from 4 min) to PM (from 23 min; see blue arrowheads) transport. (E) Flow cytometry analysis representing the mean fluorescence intensity of Tf-A647 bound at the surface of TfR-eRUSH cells. Cells were treated with biotin during indicated times and cells were subsequently switched to 4°C for Tf-A647 binding. Background fluorescence was measured by adding an acid wash step, which stripped out all surface bound Tf-A647 (grey bars). The bar graph shows the mean +/- SD of duplicates in which at least 5,000 cells per condition were acquired and is representative of 3 individual experiments. (F) Representative immunofluorescence images detecting the arrival of TfR-eRUSH at the PM. Images were acquired with spinning disk confocal microscope at indicated time points post-biotin addition. TfR-eRUSH at the PM was monitored with Transferrin coupled to Alexa Fluor 647 (Tf-A647; upper rows) or mouse anti-TfR antibody revealed with a donkey anti-mouse Alexa 647 antibody (TfR Ab; lower rows). The protein was detected at the plasma membrane starting from 20 min post-biotin addition. Scale bar = 10 µm.

From the immunoblot, it seemed that less TfR-SBP-EGFP proteins were expressed in the edited cells than the endogenous TfR from WT cells. However, quantification of the amount of proteins from bands of different sizes is not reliable due to different protein transfer efficiency. Thus, an anti-TfR antibody staining on WT and TfR-eRUSH cells was performed and the mean fluorescence intensity of the TfR staining was measured by flow cytometry. We found that TfR-eRUSH cells express less endogenous TfR than their parental cell line (Figure 1C).

Next, we carried out 3D confocal live cell imaging on TfR-eRUSH cells to determine whether TfR-eRUSH could be efficiently retained in the ER. We observed that in absence of biotin (0 min), TfR-eRUSH was retained in the ER (Figure 1D, upper panels and corresponding Movie S1). Two to six minutes after biotin addition, vesicles were released from the ER to reach the Golgi apparatus. This trend was successfully quantified by measuring the Pearson’s correlation coefficient between TfR and either Calnexin (ER) marker), GM130 (cis-Golgi) or TGN46 (trans-Golgi) at 0 min, 5 min and 15 min post-biotin addition (Figure S1C-D). While the ER released most of its vesicles, a short lag was observed at ≈ 12 minutes before observing numerous vesicles exiting from the Golgi apparatus. At 20 min, most of TfR-eRUSH was localized at the Golgi and vesicles were massively released from this location. In parallel, PM gained higher TfR-eRUSH fluorescence intensity (Figure 1D, blue arrowheads and Movie S1), indicating that the first detectable amounts of TfR-eRUSH proteins arrived at the PM at 20 min post-biotin addition.

To quantitatively measure the kinetics of TfR-eRUSH arrival at the PM, a flow cytometry assay was optimized (Figure 1E). At different times post-biotin addition, cells were incubated at 4°C to block membrane trafficking and the PM-exposed TfR was labeled using recombinant transferrin coupled to an Alexa Fluor 647 (Tf-A647) (Figure 1E). We noticed that a small fraction of TfR-eRUSH was already found at the PM even in absence of biotin (0 min), suggesting that either some aspecific Tf binding occurred or that a small amount of TfR-eRUSH was not retained by the hook. While the fluorescence signal of Tf-A647 was lowly increasing over the 20 first min after biotin addition, a three-fold increase was observed at 30 min post-biotin addition. This kinetics were confirmed by microscopy (Figure 1F) and are in agreement with our live cell imaging (Figure 1D) in which the first TfR proteins could be readily detected at the PM at ≈ 23 min post-biotin addition, then rising over time.

In conclusion, our TfR-eRUSH system represents a valid approach to study the molecular mechanism of the TfR secretory pathway in an endogenous synchronized model.

### Molecular signature of the TfR-associated membranes using TfR-eRUSH cells

To identify the molecular partners enriched in the TfR-containing membranes over time, anti-TfR affinity-purification mass spectrometry (AP-LC-MS/MS) experiments using TfR-eRUSH lysates obtained from mechanical cell disruption was performed at different time points post-biotin addition. AP-LC-MS/MS was run in quadruplicate and > 2000 proteins were identified in each sample. Differential temporal analysis identified 557 proteins enriched at T15 compared to T0 (T0-T15), while no significant protein enrichment could be measured at T30 compared to T15 (T15-T30). This absence of protein enrichment between T15 and T30 could be attributed to the lack of temporal resolution and/or the fact that multiple trafficking pathways are overlapping at these times, blurring the final picture. Parallel analyses using STRING (22) (Figure 2A), and the molecular signature database MSigDB (23) (Figure 2B and Table S1) were run on the enriched protein lists from the T0-15 differential analysis. These methods were employed to highlight protein clusters and biological processes associated to neosynthesized TfR trafficking.

**Fig 2.**
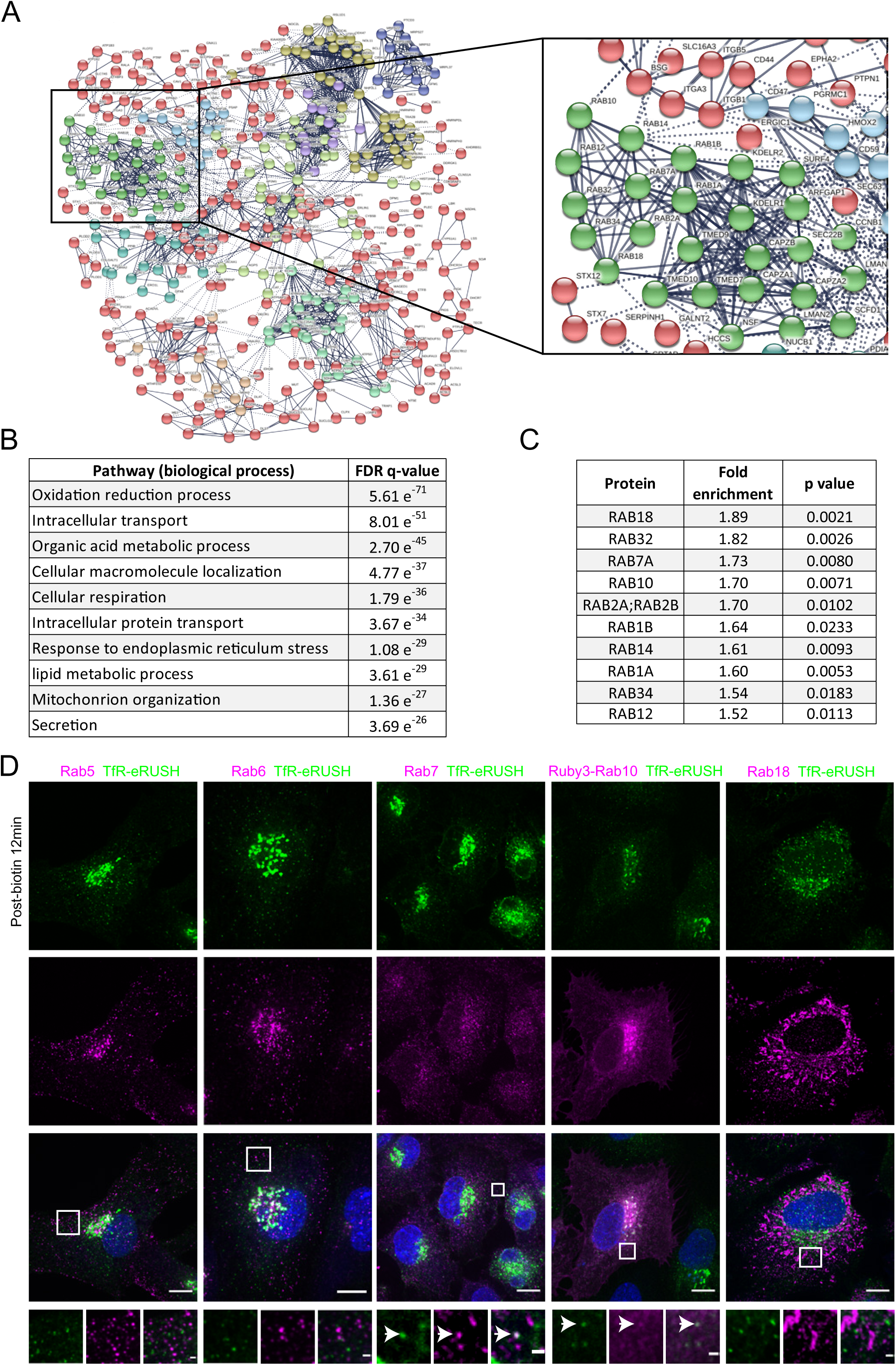
Proteomics analysis of neosynthesized TfR-containing membranes. (A-C) TfR-eRUSH cells were untreated (T0) or treated with biotin during 15 min (T15) or 30 min (T30). Mechanical cell lysis was performed at 4°C and membrane-containing TfR-eRUSH were isolated by immunoprecipitation using an anti-TfR antibody. LC-MS/MS proteomics analysis was run, and temporal protein enrichment was assessed (see material and methods for details). (A) STRING analysis shows the interaction map of the proteins that were enriched at T15 compared to T0. Color-codes highlight clusters of proteins of related functions. (B) Gene ontology of the proteins enriched at least 1.5 times with a significant p value (< 0.05) at T15 compared to T0 (T0-T15) were investigated using the online GSEA online software. Relevant GO pathways and their corresponding FDR (False-discovery rate) values are reported for each differential analysis. (C) Among the 20 Rab proteins identified by LC-MS/MS (see Table S2), the fold enrichment and p values of the ones that were significantly identified at T15 compared to T0 (T0-T15) are reported. No Rab protein were significantly enriched at other differential time points. (D) Representative confocal images from a single z-stack indicate the distribution of TfR-eRUSH treated for 12 min with biotin relative to the endogenous Rab5, Rab6, Rab7, Rab18 proteins and the exogenously expressed Ruby3-Rab10. TfR-eRUSH co-distributed with Rab7 and Ruby3-Rab10 (zoomed panel, white arrow). Scale bar = 10 µm. Zoomed regions from white squares were represented with a scale bar = 1 µm.

The pathways “intracellular transport”, “Cellular macromolecular localization”, “intracellular protein transport” and “secretion” were highly enriched compared to T0 as shown by the low false-discovery rates (FDR) values, an expected result due to the nature of the assay (Figure 2B). Moreover, the pathways associated to “exocytosis” (FDR = 4.75 10^-23^) and “Golgi vesicle transport” (FDR = 2 10^-15^) were also significantly enriched to a lower extend. As a proof of concept, we confirmed that TMED10, a protein identified as enriched in our proteomics analyses was indeed recruited to TfR secretory vesicles (Table S1, Figure S2A). TMED10 is involved in the COPII vesicle-mediated anterograde transport (24) and incorporated in a subset of extracellular vesicles (25), and thus we could confirm the relevance of our differential proteomics approach.

The pathways “oxidation reduction process”, “cellular respiration” and “mitochondrion organization” also scored significant low FDR values. ER and mitochondrial membranes are well-known to tightly interact (26) and recent work proposed that endosome-mitochondria interactions are important for the release of iron (27). Here, the mitochondria-associated proteins identified may be the result of association of distinct membranes during the immunoprecipitation rather than actual presence of TfR within mitochondria. In fact, proximity was observed between TfR-eRUSH and mitotracker-labeled mitochondria (Figure S2B). By live cell imaging, we visualized some rare events of mitochondria “associated” with vesicles containing TfR-eRUSH that seemed to bud off the ER, but the resolution achieved with our spinning disk confocal microscope does not allow us to draw significant conclusion (Figure S2C and Movie S2).

Proteins regulating intracellular trafficking may be differentially recruited on vesicular membranes to activate a specific trafficking route. Therefore, we chose to further investigate the role of Rab proteins as they are well-known small GTPase regulators of intracellular membrane traffic. In our AP-LC-MS/MS dataset, we detected a total of 20 Rab proteins (Table S2). No Rab proteins were enriched in TfR-containing membranes at T15-T30, but 10 Rab proteins were significantly enriched at T0-T15 with a fold change above 1.5 times (Figure 2C). Rab1A, Rab1B and Rab18 were significantly enriched at T15 compared to T0, an expected result as these Rabs regulates vesicle trafficking between the ER and the cis-Golgi (28, 29), Rab18 being also found on a subset of extracellular vesicles (25). Rab10, Rab14 and Rab6A were also enriched at T15 compared to T0, although Rab6 did not reach significance (Table S2). These Rabs have previously been involved in post-Golgi trafficking (13,30,31), further indicating that our approach is relevant to identify molecular partners involved in the secretory pathway. The Rab12 and Rab34 proteins were also identified, but their function has not been extensively studied. Yet they both may play a role in protein degradation (32, 33). Finally, Rab7A, a protein usually recruited at the limiting membrane of late endosomes that can serve as degradation signal (34), was significantly enriched at T15 compared to T0. Rab7A showed one of the highest fold enrichment score and the greatest number of unique peptides identified by LC-MS/MS (Figure 2C and Table S2), an intriguing result that we aimed to explore thereafter.

To further investigate the relevance of the Rabs identified in our proteomics analyses, the distribution of Rab5, Rab6, Rab7, Ruby3-Rab10 and Rab18 was imaged in TfR-eRUSH cells treated for 12 min with biotin (Figure 2D). While no colocalization was observed between TfR-eRUSH and Rab5 nor Rab18, association with Rab6, Rab7, Rab10 was seen.

### Rab7 is significantly enriched onto post-Golgi TfR-eRUSH vesicles

To further characterize the recruitment of Rab7A on TfR-containing secretory vesicles, we performed live cell imaging by spinning disk confocal microscopy on TfR-eRUSH cells transfected with a Ruby3-Rab7A construct under the control of the weak promoter L30 (to minimize overexpression). Starting from 7 min post-biotin addition, we noticed the presence of post-TGN TfR-eRUSH signal associated to Rab7A positive vesicles (Figure 3A and Movie S3). To better appreciate whether TfR-eRUSH and Rab7A were found on the same vesicles (as opposed to two distinct vesicles in close proximity), we artificially swollen these compartments using Apilimod, a PIKfyve inhibitor (35), and indeed, we could identify that TfR-eRUSH-positive vesicles were decorated with Rab7A at their limiting membrane (Figure 3B). These data were reminiscent of a recent work nicely demonstrating that post-Golgi vesicles were positive for Rab6 (13) and indeed in our model, TfR-eRUSH was also trafficking through Rab6 (Figure S3A).

**Fig 3.**
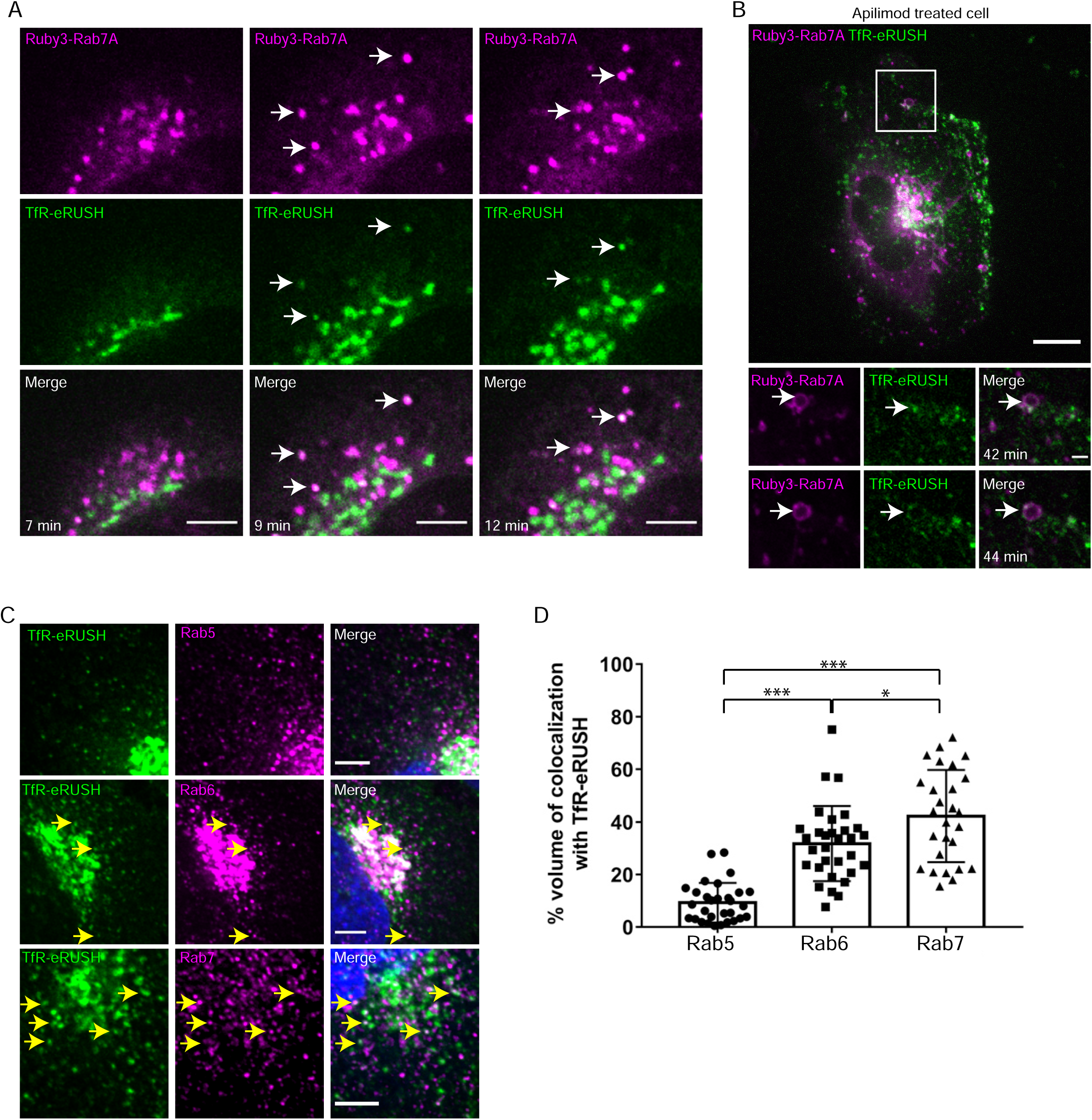
Identification of Rab7 as an intermediate compartment of neosynthesized TfR trafficking. (A) Live cell imaging using a spinning disk confocal microscope shows localization of TfR-eRUSH in Ruby3-Rab7A transfected cells. Representative images were extracted from a single-plane and TfR-eRUSH (green) was visualized within Ruby3-Rab7A (magenta) positive vesicles (white arrows) at 7 min, 9 min and 12 min post-biotin addition, scale bar = 5 µm. (B) TfR-eRUSH cells expressing Ruby3-Rab7A were imaged as in A after treatment with 40 nM Apillimod for 30 min to increase the size of Ruby3-Rab7A vesicles. Biotin was added to release TfR-eRUSH and representative images performed with the spinning disk microscope were extracted as a single-plane at 42- and 44-min post-biotin addition. TfR-eRUSH (green) localizes at the limiting membrane of Ruby3-Rab7A vesicles (magenta) (white square, scale bar = 10 µm). The zoomed regions from the white square highlight single and merge staining of the Ruby3-Rab7A containing TfR-eRUSH (scale bar = 1 µm). The white arrows indicate the repartition of TfR-eRUSH at the limiting membrane of the Ruby3-Rab7A vesicle. Of note, trafficking kinetics were much longer following Apilimod treatment. (C-D) TfR-eRUSH cell were treated 15 min with biotin, fixed and stained for endogenous Rab5, Rab6 or Rab7 using specific rabbit antibodies revealed by a donkey anti rabbit Alexa Fluor 647 antibody. (C) Representative images from a single z-stack indicate the localization of TfR-eRUSH relative to the endogenous Rab5, Rab6, Rab7 using spinning disk microscope. TfR-eRUSH colocalization with Rab6, Rab7 is represented with yellow arrows. Scale bar = 5 µm. (D) The graph represents the quantification of the volume of TfR-eRUSH (+/- SEM) colocalizing with Rab5, Rab6 or Rab7. Data represents n = 30 cells (Rab5), n = 31 cells (Rab6), n = 27 cells (Rab7) per condition from 3 independent experiments and student t-test was run to determine significance (* p value < 0.05 and *** p value < 0.001).

Quantification of TfR-eRUSH association with indicated Rabs was then performed using antibody staining on TfR-eRUSH cells fixed at 15 min post-biotin addition. As expected, the percentage of non-Golgi TfR-eRUSH signal associated to Rab5 was very low (9.3% +/- 1.3), while association with Rab6 and Rab7 was relatively high (31.7% +/- 2.5 and 42.3% +/- 3.3, respectively; Figure 3C-D). Interestingly however, TfR-eRUSH vesicles would harbor either Rab7A or Rab6, but no post-Golgi TfR-eRUSH-Rab6-Rab7A triple colocalization was seen (Figure S3B).

Together, our proteomics analysis revealed that several Rabs are enriched onto secretory TfR-containing vesicles and that Rab7A represent an unexpected protein recruited in the neosynthetic secretory pathway.

### Neosynthesized TfR associates with non-degradative Rab7 vesicles

Rab7 is known to direct late endosomal compartments toward degradative Lamp1-positive compartments (36). By immunostaining, we observed that a subset of TfR-eRUSH was Rab7-positive and Lamp1-negative (Figure 4A, yellow arrow), but we could also see triple colocalization of TfR-eRUSH, Rab7, and Lamp1 (Figure 4A, white arrowheads). However, mapping the association with Lamp1 is not sufficient to define lysosomal compartments since a recent study demonstrated that TfR is co-sorted with Lamp1 into post-TGN secretory vesicles *en route* to the PM (37). Moreover, Lamp1 was also identified in our proteomics analysis (Table S1).

**Fig 4.**
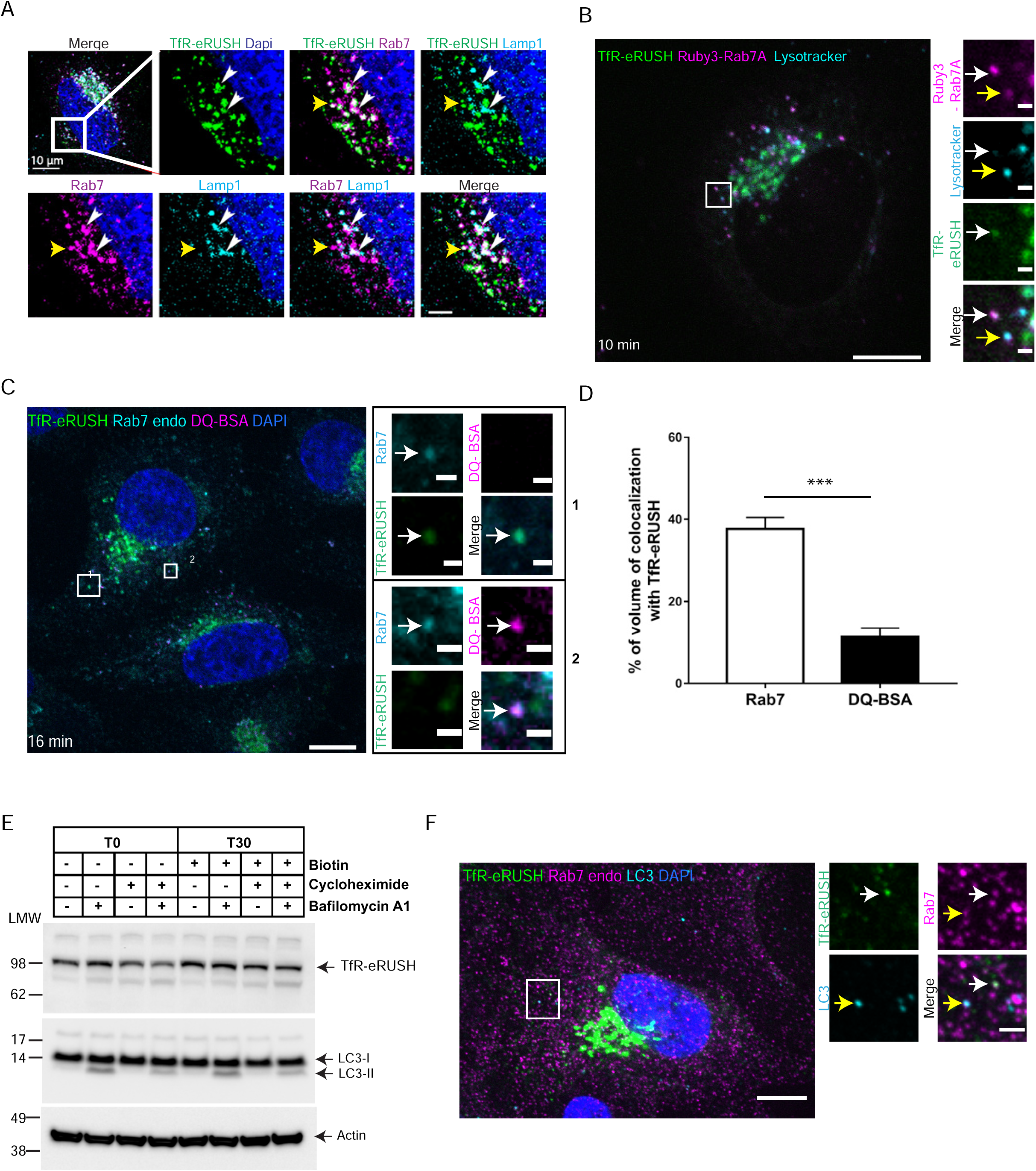
TfR-eRUSH transiting through Rab7 vesicles, localizes at the plasma membrane. (A) Representative images from a single z-stack indicate TfR-eRUSH co-distributed with Lamp1 and Rab7 positive vesicles. TfR-eRUSH cells were treated with biotin for 7 min, fixed and stained using a rabbit anti-Rab7 antibody and a mouse anti-Lamp1 antibody revealed by a donkey anti-rabbit coupled to Alexa Fluor 568 and a donkey anti-mouse coupled to Alexa Fluor 647. The images are single-plane crops from the white square of the upper left image. The snapshots show dual- and triple staining and the merge (lower right panel). TfR-eRUSH (green) co-distributed with Rab7 (magenta) and Lamp1 (cyan) in perinuclear localized structures (white arrowheads). Some Rab7 positive vesicles contain TfR-eRUSH but no Lamp1 (yellow arrows). (B) Live cell imaging indicates that Rab7A positive vesicles containing TfR-eRUSH are not labeled by lysotracker. Cells were transfected with Ruby3-Rab7A for 24h. Before imaging, cells were incubated for 30 min with lysotracker (50 nM) then biotin was added to induce the release of TfR-eRUSH. Images extracted at 10 min post-biotin addition, were acquired with spinning disk confocal microscopy and represented as a single plane (scale bar = 10 µm). White squares showed the zoomed panels with single staining and the merge. The white arrow indicates that Ruby3-Rab7A (magenta) co-distribute with TfR-eRUSH (green) while the yellow arrow shows that lysotracker (cyan) co-distribute with Ruby3-Rab7A (magenta). Scale bar= 2 µm. (C) Immunofluorescence images indicate that Rab7A vesicles containing TfR-eRUSH does not contain DQ-BSA. TfR-eRUSH cells were incubated with 10 µg/ml of DQ-BSA for 6 h in the presence of 1 µg/ml avidin. Then, TfR-eRUSH cells were treated with biotin for 16 min, fixed and stained with rabbit anti-Rab7 antibody. Secondary antibody donkey anti rabbit Alexa Fluor 647 was used. Images were acquired with spinning disk confocal microscopy and are represented as a single plane (scale bar = 10 µm). White squares indicated zoomed regions with single staining and the merge. TfR-eRUSH (green) co-distributed with Rab7 (cyan) (square 1) and DQ-BSA (magenta) co-distributed with Rab7 (cyan) (square 2). Scale bar = 2 µm. (D) The bar graph represents the quantification of the TfR-eRUSH vesicles colocalizing with Rab7A or DQ-BSA. Data represents n = 32 cells (DQ-BSA) and n = 28 cells (Rab7) from 3 independent experiments (+/- SEM). Student t-test was run to determine significance (*** p value < 0.001). (E) Western blot analysis indicate that TfR-eRUSH is not degraded following biotin addition. TfR-eRUSH cells were incubated for 4 h in the presence or absence of 50 µg/ml cycloheximide and 100 nM bafilomycin A1 as indicated. Biotin was added for 0 or 30 min and cells were lysed for western blot analysis. Actin was used as a loading control. The presence of LC3-II over LC3-I confirmed the inhibitory effect of bafilomycin A1 on protein degradation. Low molecular weight protein marker (LMW) was used for molecular weight estimation. (F) Immunofluorescence images indicates that LC3 does not colocalize with Rab7 and TfR-eRUSH. Cells were treated with biotin for 15 min, fixed and stained with rabbit anti-Rab7 antibody, and mouse anti-LC3 antibody. Secondary antibody donkey anti rabbit Alexa Fluor 647 and donkey anti-mouse Alexa 561 were used. Images were acquired with spinning disk confocal microscopy and are represented as a single plane (scale bar = 10 µm). White squares indicated zoomed regions with single staining and the merge. TfR-eRUSH (green) co-distributed with Rab7 (magenta) but not with LC3 (cyan) Scale bar = 2 µm.

Therefore, to better address whether the TfR-Rab7 vesicles correspond to degradative compartments, pH acidity and proteolytic activity was measured (Figure 4B-C). TfR-eRUSH cells were transfected with Ruby3-Rab7A and were visualized by live imaging at 10 min post-biotin addition. Lysotracker was used as a readout for relative pH acidity (a brighter signal corresponding to a lower pH). Our data shows that TfR-eRUSH vesicles harboring Rab7A had little-to-no lysotracker signal (Figure 4B, white arrow), indicating that these vesicles do not exhibit features of classical proteolytic compartments. To assess for actual degradative properties of these vesicles, we pre-incubated the cells with DQ-BSA, a bovine serum albumin (BSA) protein that contains self-quenched fluorescent dyes that fluoresce only when the BSA is cleaved, and stained the cells with an anti-Rab7 antibody (Figure 4C). Quantification of the percentage of TfR-eRUSH colocalizing with Rab7 or DQ-BSA demonstrated that the TfR were mainly found in Rab7 vesicles devoid of degraded DQ-BSA (Figure 4D). Finally, because the TfR was engineered to incorporate SBP and EGFP, we checked whether a significant subset of protein was sent for degradation. However, cells treated with Bafilomycin A1 (to prevent protein degradation) did not induce an accumulation of TfR-eRUSH, while it induced LC3-II accumulation as expected (Figure 4E). To make sure that the absence of visible degradation was not due to neosynthesized TfR-eRUSH replenishment, cells were co-treated with Bafilomycin A1 and cycloheximide, a translation inhibitor. In this context, we could not observe any accumulation of TfR (Figure 4E) and we were not able to detect degradation products using either anti-TfR or anti-EGFP antibodies (Figure S4), suggesting that TfR-eRUSH is not significantly sent for degradation. Of note, this experiment also indicates that the induction of the eRUSH by the addition of biotin is not accompanied by the induction of autophagy as LC3-II is not upregulated (Figure 4E). To further confirm this, LC3 staining was performed and we could not observe any LC3 staining colocalizing with Rab7-positive TfR-eRUSH vesicles (Figure 4F), indicating that they do not correspond to autophagosomes.

### Rab7A-vesicles are intermediate compartments mediating the transport of a subset of neo-synthesized TfR-eRUSH to the PM

To correlate the enrichment over time of Rab proteins to a biological function, we next carried out a siRNA-based screen targeting 12 members of the Rab protein family. Silencing of 12 Rabs and a non-relevant target was performed using a pool of 4 siRNA per target in two independent experiments (Figure S5A). The amount of TfR-eRUSH at the PM was measured by flow cytometry as in Figure 1E, and fold enrichment of T15 over T0 (Figure 5A) was determined. As Rabs may affect other cellular processes, the amount of TfR-eRUSH at steady-state was measured by flow cytometry (Figure S5).

**Figure 5.**
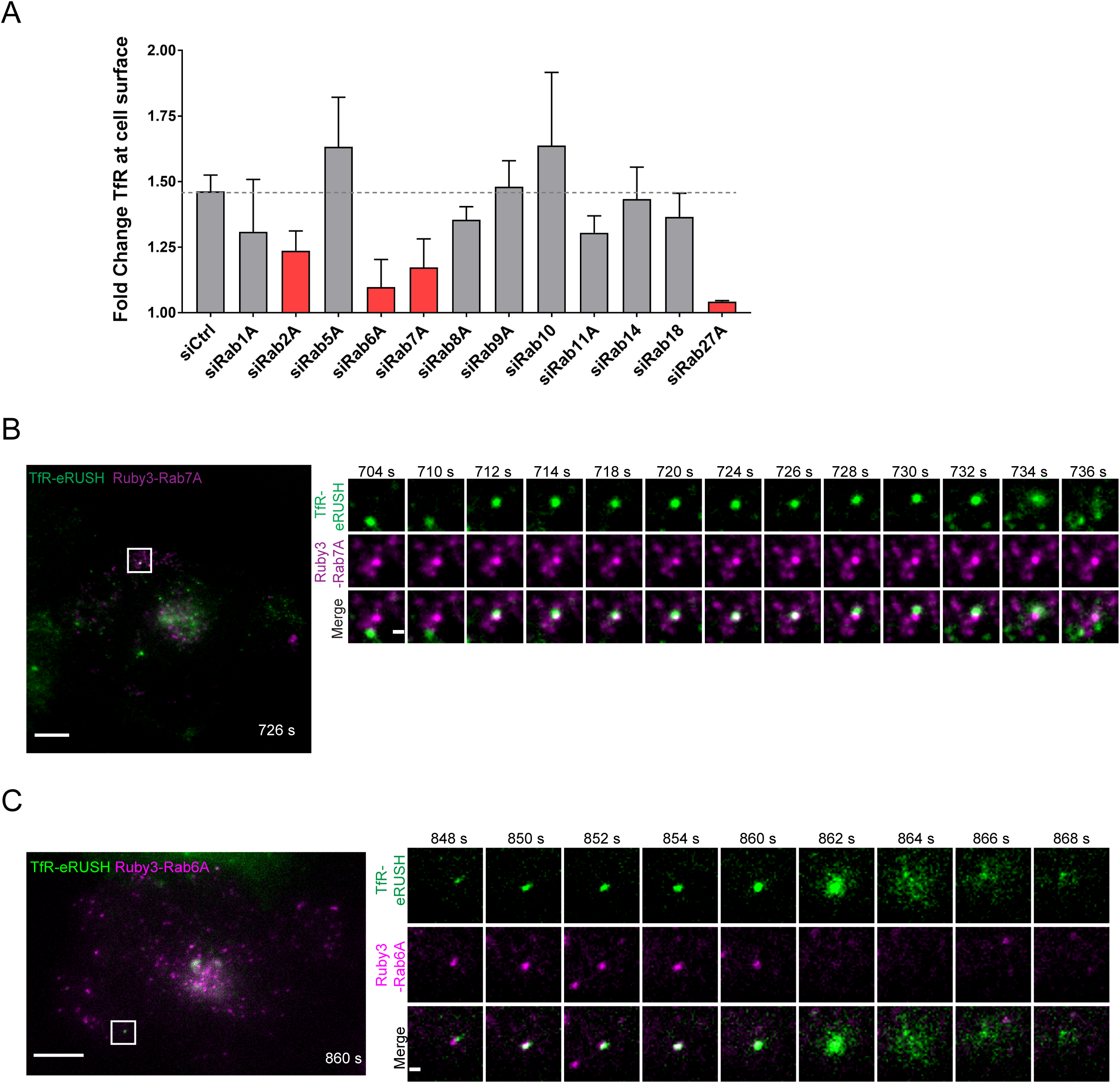
Rab7A is involved in transport of TfR-eRUSH at the plasma membrane. (A) TfR-eRUSH cells were treated with siRNA sequences targeting 12 different Rab mRNAs and a non-targeting siRNA control. After 48 h post-transfection, cells were untreated (T0) or treated with biotin for 15 min (T15). Measure of the amount of TfR-eRUSH at the PM was performed by flow cytometry as in Fig 1E. The bar graph represents the mean fold change +/- SD corresponding to the ratio between the Tf-A647 MFI measured at 15 min and 0 min from two individual experiments performed in duplicates in which at least 2,000 cells were analyzed. Anova and Student t-test were run to assess for significance (red bar graph: p value < 0.05 for siRab2A, siRab6A and siRab7A and p value < 0.001 for siRab27A). (B) TfR-eRUSH cells transfected with Ruby3-Rab7A were imaged by TIRF microscopy 24 h post-transfection. Cells were imaged from 5- to 25-min post-biotin addition. A representative image extracted from the Movie S4 was shown at 726 s (scale bar = 10 µm). The white square indicates the cropped region represented in the right panels. In these panels, a Ruby3-Rab7A (magenta) vesicle carrying TfR-eRUSH (green) was tracked from 704 s to 736 s. Scale bar = 1 µm. (C) TfR-eRUSH cells transfected with Ruby3-Rab6A were imaged by TIRF microscopy 24 h post-transfection. Cells were imaged from 5- to 25-min post-biotin addition. A representative image extracted from movie S5 was shown at 860 s (scale bar = 10 µm). The white square indicates the cropped region represented in the right panels. In the right panels a Ruby3-Rab6A (magenta) vesicle carrying TfR-eRUSH (green) was tracked from 848 s to 868 s. It indicates a Rab6A vesicle releasing a TfR-eRUSH directly to the PM. Scale bar = 1 µm.

At T0-T15, silencing of Rab27A or Rab6A showed significant decrease of PM-associated TfR (Figure 5A) compared to the non-relevant siRNA control. These findings were in agreement with the role of these Rabs in protein secretion (13,38,39), validating our approach. In contrast, Rab10 silencing had no detectable effect on TfR trafficking to the PM, while we found it enriched in our proteomics analysis (Figure 2C). Interestingly however, silencing of Rab7A showed a significant inhibition of TfR-eRUSH arrival at the PM at T0-15. Although Rab7A is known for its role in endocytic retrograde trafficking to late endosomes and lysosomes (34), this is consistent with our AP-LC-MS/MS data (Figure 2C), further indicating that Rab7 could participate in the transport of post-TGN TfR vesicles.

To directly determine the fate of the Rab-harboring post-Golgi TfR-eRUSH vesicles, total internal reflection fluorescence (TIRF) microscopy was performed on cells transfected with Ruby3-Rab7A (Figure 5B and Movie S4) or Ruby3-Rab6A (Figure 5C and Movie S5). After 12 min post-biotin addition, the arrival of TfR-eRUSH was observed in the evanescent TIRF field. We monitored events during which Rab7-positive vesicles became positive for TfR-eRUSH for several seconds (Figure 5B; from 704 s to 736 s) before the two signals segregated again, followed by a TfR-eRUSH signal burst, indicative of PM fusion (734 s). In sharp contrast, Ruby3-Rab6A remained associated to TfR-eRUSH vesicles until fusion occurred (Figure 5C; 848-868 s).

These observations indicated that Rab7A vesicles are used as intermediate compartments in TfR trafficking after its release from the TGN. Unlike Rab6A, Rab7A vesicles do not accompany neosynthesized TfR all the way to the PM and thus, other partners are likely involved downstream of the Rab7-TfR vesicle trafficking.

## Discussion

Description of the different pathways mediating transport of neo-synthesized receptors to the PM has been studied for decades. Being able to specifically observe the anterograde pathway has always been a challenge as its visualization overlaps with other trafficking routes, including the overrepresented endocytosis and recycling pathways. To visualize protein transport under physiological conditions, we combined the RUSH system to the CRISPR/Cas9 technology. Using TfR as a model, we generated a stable cell line expressing endogenous levels of the receptor fused to EGFP and the SBP tag required for the RUSH system. TfR function and trafficking are well described but the partners involved in neosynthesized TfR trafficking to the PM are not well characterized. The eRUSH (edited-RUSH) approach was coupled to quantitative proteomics experiments and cytometry-based screening to identify the molecular partners involved in the neosynthetic pathway of the TfR. Unexpectedly, we observed that a significant subset of TfR transits through Rab7-positive vesicles during its trafficking to the PM.

The trafficking kinetics of neosynthesized TfR-eRUSH was similar to the overexpressed TfR in the RUSH system which was previously described to reach the PM ≈ 30 min post-biotin addition (37). The advantage of our eRUSH is that no or minimal amount of “ghost” untagged proteins are expressed in TfR-eRUSH, allowing quantitative single molecule counting as well as whole TfR functional analysis. Moreover, the eRUSH also represents a powerful knock-away system, similar to other methods (40), but without the problem of competition with the wild type version of the protein co-expressed in the cell. In contrast to classic cDNA transfection, CRISPR/Cas9-based gene editing of TfR allows the conservation of the regulatory genomic environment of the gene. This parameter is particularly important for proteins such as TfR as its transcriptional/translational regulation is a finely regulated process (41, 42).

Using AP-LC-MS/MS, we could track the local TfR environment at different times post-biotin addition and identify proteins co-distributing with TfR-extracted membranes. Whereas previous siRNA-based screens studying the secretory pathway allowed the identification of important novel partners (43), our eRUSH-based proteomics is based on a non-interfering approach, and thus, it provides new complementary information to previous studies. Pathway analysis revealed relevant enriched biological processes as well as less expected ones. Indeed, an enriched proportion of mitochondrial proteins at 15 min post-biotin addition was observed. We propose that this result is due to ER-mitochondria membrane contacts sites and may not be relevant to the biosynthetic pathway of TfR.

Some Rabs can act together in the exocytosis process, such as Rab6 and Rab8 (44) or Rab3 and Rab27 (45). We detected > 35% of TfR-eRUSH-containing vesicles harboring a Rab6-positive staining (Figure 3D), but our data suggest however that Rab6 and Rab7 do not intervene at the same stage of the secretory pathway and/or in the same type of vesicular transport, as shown by the absence of overlap between the Rab6 and Rab7 staining (Figure S3B). Moreover, by TIRF microscopy (Figure 5B-C), we noticed two different processes of TfR transport using Rab6A or Rab7A, further indicating that these two Rabs likely correspond to two distinct secretory routes.

Combining results from the AP-MS/MS and siRNA screen, only one Rab was significantly standing out: Rab7. Rab7 is mostly known to mediate cargo trafficking between late endosomes and lysosomes (34) and it was unexpected to find it involved in the neosynthetic pathway. By electron microscopy, a group observed the presence of neosynthesized TfR inside endosome-like structures (6). Moreover, it was demonstrated that Rab7 was not involved in recycling of TfR at the PM as depletion of Rab7A had no effect on TfR re-localization to the PM (34) and thus, it is unlikely that our observations would be the result of marginal PM-associated TfR endocytosis at early times post-biotin addition.

A legitimate thought is to believe that the post-golgi Rab7-decorated TfR-eRUSH vesicles correspond to a degradative pathway. However, extensive analyses of these vesicles clearly show that they stain mostly negative/dim to lysotracker, they are not proteolytically active (DQ-BSA marker) nor autophagosomes, while autophagy is not induced by the biotin treatment (Figure 4). Moreover, the full membranes of the western blot analysis show no degradative product, further demonstrating that TfR-eRUSH vesicles harboring Rab7 are not degradative and actually, direct evidence support PM-targeting of these vesicles (Figure 5B). Yet, the function of these vesicles as compared to the Rab6-positive vesicles, remain to be determined.

A recent study by Chen and colleagues indicated that neosynthesized TfR was sorted out with the Lamp1 protein in vesicles exiting the TGN (37). These vesicles were devoid of the mannose-6P receptor (M6PR), which was used as a marker for Golgi-to-endosome route (46). In our hands, we found that M6PR was absent of the TfR-eRUSH vesicles harboring Rab7 (not shown). They concluded that the TfR^+^ Lamp1^+^ vesicles were *bona fide* secretory vesicles *en route* to the PM. In our study, a subset of vesicles containing TfR-eRUSH and Lamp1 were also decorated by Rab7 at time points corresponding to TGN exit. These vesicles may correlate with the ones described by Chen *et al.* but their comprehensive composition and function in the secretory pathway remains to be fully determined.

We suggest that Rab7 could act as an intermediate compartment for neosynthesized TfR transport. Although the role of Rab7 on these secretory vesicles remains to be determined, one could hypothesize that Rab7 regulates the trafficking of cargos with specific post-translationally modifications. Alternatively, this pathway could transport cargos dedicated to specific PM domains. Recently, Rab7 has been mapped not only to late endosomes and lysosomes but also at the ER, TGN and mitochondrial membranes, a localization maintained by the retromer complex (47), and thus, it is likely that Rab7 exerts pleiotropic roles.

## Materials and Method

### Cell culture

SUM159 cells were cultured in DMEM/F-12 GlutaMAX (GIBCO), supplemented with 5% fetal bovine serum (FBS; Dominique Dutscher), 500 µg/ml penicillin-streptomycin (GIBCO), 1 µg/ml hydrocortisone (Sigma-Aldrich), 5 µg/ml insulin (Sigma-Aldrich), and 10 mM HEPES (GIBCO) (complete medium). Cells were maintained at 37°C and 5% CO_2_.

### Generation of the TfR-eRUSH CRISPR/Cas9 edited cell line

Gene editing of the SUM159 cells to fuse the GGSGGSGGS spacer, the SBP and EGFP sequences to the C-ter of TfR a CRISPR/Cas9 strategy was used as previously described (21, 48). Briefly, three genetic tools were co-transfected using the transfection reagent TransfeX (ATCC): 1) a plasmid coding for CRISPR-associated protein 9 (Cas9), a template plasmid; 2) a linear PCR product used to transcribe the tracrRNA and guide RNA (gRNA) targeting ATAGCTTCCATGAGAACAGC (corresponding to a region near the genomic TfR stop codon) under the control of the human U6 promoter; 3) a donor DNA construct (serving as template during homologous recombination) corresponding to the spacer, SBP and EGFP sequences flanked by ≈ 800 bp upstream and 800 bp downstream of the TfR stop codon. Single cell sorting of EGFP-positive cells was performed and homo/heterozygotic monoclonal cell lines expressing endogenous TfR-eRUSH were screened by PCR using the forward primer 5’ CTCACACGCTGCCAGCTTTA 3’ and reverse primer 5’ TTCAGCAGAGACCAGCCCTT 3’.

A clone that was edited on both alleles was further transduced with a lentiviral vector coding the puromycine resistance gene and for the “hook” consisting of the streptavidin protein linked to the KDEL motif (11). Upon puromycin selection, the SUM159 TfR-eRUSH cells were expanded and stocks for the original tube were maintained in liquid nitrogen.

### Plasmids

The Ruby3-Rab7A, Flag-Apex-Rab7A, Ruby3-Rab6A and Ruby3-Rab10 cDNA constructs cloned into pBS vectors under the control of the weak promoter L30, were generated by the Montpellier Genomics Collections (MGC).

### Antibodies and reagents

For immunofluorescence, primary antibodies used were mouse anti-GM130 (1/1000; BD bioscience), sheep anti-TGN46 (1/1000, Bio-Rad), rabbit anti-Calnexin (1/1000, Elabscience), mouse anti-Lamp1 (1/100, BD bioscience), rabbit anti-Rab7 (1/250, Cell Signaling Technology), rabbit anti-Rab5 (1/1000, Cell Signaling Technology), rabbit anti-Rab6 (1/1000, Cell Signaling Technology), rabbit anti-Rab18 (1/200, Sigma-Aldrich), rabbit anti-TMED10 (1/500, Sigma-Aldrich), mouse anti-LC3 (1/1000, Sigma-Aldrich) and mouse anti-TfR (1/250, Miltenyi Biotec). Secondary antibodies used were Alexa fluor 568 donkey anti-sheep (1/1000, life technologies), Alexa fluor 568 donkey anti-rabbit (1/1000, Thermo Fisher Scientific), Alexa fluor 647 donkey anti-mouse (1/1000, Thermo Fisher Scientific). Antibodies used for immunoblotting were rabbit anti-TfR (1/1000, Aviva Systems Biology), mouse anti-beta actin (abcam), mouse anti-GFP (1/1000, Sigma-Aldrich) and rabbit anti-LC3 (1/1000, Sigma-Aldrich). Secondary antibodies used for immunoblotting were Goat anti-mouse IgG HRP antibody (1/10000, Jackson ImmunoResearch), Goat anti-rabbit IgG HRP antibody (1/10000, Jackson ImmunoResearch). Probes used for immunofluorescence were membrane-permeable MitoTracker Orange CM-H_2_TMRos (Molecular probes) used at 100 nM to label mitochondria, Lysotracker red (Life Technologies) for acidic compartments used 30 min at 50 nM, DQ-Red BSA (Life Technologies) used at 10 µg/ml in complete medium, and DAPI (1/1000, Sigma-Aldrich) used to stain the nucleus. For flow cytometry Transferrin coupled to Alexa fluor 647 (molecular probes) was used at 10 µg/ml, anti-mouse TfR (5 µg/ml, Miltenyi Biotec), and IgG mouse used as an isotype control. For deglycosylation, endoglycosidase H was used (NEB).

### Cytometry-based RUSH assay

To detect PM-localized TfR, 40 000 SUM159 cells were plated in 48 well plates and incubated in complete medium containing 0,28 μg/ml avidin (Sigma-Aldrich) for 48 hours. To initiate TfR release, cells were incubated in a fresh complete medium containing 40 μM biotin (Sigma) for the indicated amount of time at 37°C and 5% CO_2_. Then, cells were placed on ice, the media was replaced with ice-cold PBS and cells were maintained at 4°C for 15 min. Cells were incubated for 20 min with 10 µg/ml of Tf coupled to an Alexa Fluor 647 (Tf-A647; Molecular probes) diluted in PBS pH 7.0 at 4°C. Unbound Tf-A647 was washed two times with cold PBS and cells were detached with 5 mM EDTA. Cells were collected and centrifuged at 400 g for 15 min at 4°C. Cell fixation was carried out with 4% paraformaldehyde (PFA) for 20 min at room temperature and after three washes, they were resuspended in a flow cytometry buffer (PBS pH7.0, 0.5% BSA, 0.5 mM EDTA). Samples were run on a Cytoflex flow cytometer (Beckman Coulter) equipped with 488 and 640 nm lasers and 4 filter set.

### SiRNA screen

A pool of four different siRNAs for each of the 12 selected Rab proteins and a non-targeting siRNA control was purchased as a custom-made siGenome Smart pool cherry-pick library (Dharmacon, Horizon Discovery; see details in Table S3). Forty thousand SUM159 cells were seeded in 48-well plates and on the next day, 3 pmol of siRNA were transfected using lipofectamine 2000 (ThermoFisher Scientific) according to the manufacturer’s instructions. Cells were further incubated 48 h in complete medium in presence of 0.28 μg/ml of avidin. The day of the experiment, cells were incubated at different time points with 40 μM of biotin. The cytometry-based assay for PM-localized TfR described above was used for sample analysis.

### Immunoprecipitation of TfR-eRUSH

For immunoprecipitation of the TfR-eRUSH proteins, SUM159 cells were plated in 20 mm sterile culture-treated petri dishes (Corning) for 48 h in complete medium with 0.28 μg /ml of avidin. Upon TfR-eRUSH release by addition of 40 µM biotin, the cells were incubated on ice, washed with ice-cold PBS and scraped into 1 ml of ice-cold isolation buffer (PBS devoid of Ca^2+^ and Mg^2+^, 0.1% BSA, 2 mM EDTA, pH7.4). Cells were lysed at 4°C by mechanical lysis using a 22G needle and the resulting lysate was centrifuged at 2 000 g for 15 min at 4°C. Supernatants were incubated for 2 h at 4°C with 2 μg of anti-TfR antibody previously coupled to Dynabeads (Thermo Fisher Scientific). The immunoprecipitated TfR-eRUSH-containing membrane fractions were washed five times with ice-cold PBS at 4°C before elution.

### Mass spectrometry-based quantitative proteomics

#### Sample preparation

The immunoprecipitated samples were resuspended in Laemmli buffer and the antibody-conjugated magnetic beads were removed. Protein concentration was determined using the RC-DC protein assay (Bio-Rad) according to the manufacturer’s instructions and a standard curve was established using BSA. For each sample, 8 µg of protein lysate was concentrated on a stacking gel by electrophoresis. The gel bands were cut, washed with ammonium hydrogen carbonate and acetonitrile, reduced and alkylated before trypsin digestion (Promega). The generated peptides were extracted with 60% acetonitrile in 0.1% formic acid followed by a second extraction with 100% acetonitrile. Acetonitrile was evaporated under vacuum and the peptides were resuspended in 16 µL of H20 and 0.1% formic acid before nanoLC-MS/MS analysis.

#### NanoLC-MS/MS analysis

NanoLC-MS/MS analyses were performed on a nanoACQUITY Ultra-Performance LC-system (Waters, Milford, MA) coupled to a Q-Exactive Plus Orbitrap mass spectrometer (ThermoFisher Scientific) equipped with a nanoelectrospray ion source. Samples were loaded into a Symmetry C18 precolumn (0.18 x 20 mm, 5 μm particle size; Waters) over 3 min in 1% solvent B (0.1% FA in acetonitrile) at a flow rate of 5 μL/min followed by reverse-phase separation (ACQUITY UPLC BEH130 C18, 200 mm x 75 μm id, 1.7 μm particle size; Waters) using a binary gradient ranging from 1% and 35% of solvent A (0.1 % FA in H2O) and solvent B at a flow rate of 450 nL/min. The mass spectrometer was operated in data-dependent acquisition mode by automatically switching between full MS and consecutive MS/MS acquisitions. Survey full scan MS spectra (mass range 300-1800) were acquired in the Orbitrap at a resolution of 70K at 200 m/z with an automatic gain control (AGC) fixed at 3.10^6^ ions and a maximal injection time set to 50 ms. The ten most intense peptide ions in each survey scan with a charge state ≥2 were selected for MS/MS. MS/MS spectra were acquired at a resolution of 17,5K at 200 m/z, with a fixed first mass at 100 m/z, AGC was set to 1.10^5^, and the maximal injection time was set to 100 ms. Peptides were fragmented in the HCD cell by higher-energy collisional dissociation with a normalized collision energy set to 27. Peaks selected for fragmentation were automatically included in a dynamic exclusion list for 60 s. All samples were injected using a randomized and blocked injection sequence (one biological replicate of each group plus pool in each block). To minimize carry-over, a solvent blank injection was performed after each sample. A sample pool comprising equal amounts of all protein extracts was constituted and regularly injected 4 times during the course of the experiment, as an additional quality control (QC). Protein identification rates and coefficients of variation (CV) monitoring of this QC sample revealed very good stability of the system: 2207 of the 2271 identified proteins, namely 97%, showed a CV value lower than 20% considering all 4 injections.

#### Data interpretation

Raw MS data processing was performed using MaxQuant software (v 1.5.8.3 (49)). Peak lists were searched against a composite database including all *Homo sapiens* protein sequences extracted from UniprotKB-SwissProt (version April 2019; taxonomy ID: 9606) using the MSDA software suite (50). MaxQuant parameters were set as follows: MS tolerance set to 20 ppm for the first search and 5 ppm for the main search, MS/MS tolerance set to 40 ppm, maximum number of missed cleavages set to 1, Carbamidomethyl (C) set as fixed modification, Acetyl (Protein N-term) and Oxidation (M) set as variable modifications. False discovery rates (FDR) were estimated based on the number of hits after searching a reverse database and was set to 5% for both peptide spectrum matches (minimum length of seven amino acids) and proteins. Data normalization and protein quantification was performed using the LFQ (label free quantification) option implemented in MaxQuant (49) using a “minimal ratio count” of two. The “Match between runs” option was enabled using a 2 min time window after retention time alignment. All other MaxQuant parameters were set as default.

To be validated, proteins must be identified in all four replicates of one condition at least. The imputation of the missing values and differential data analysis were performed using the open-source ProStaR software (51). Two runs of imputation were applied, the “SLSA” mode was applied for the POV (partially observed values) and the “del quantile” for the MEC (missing in the entire condition). Pairwise comparisons were performed using a Limma t-test on protein intensities. P-values calibration was performed using the pounds calibration method and the FDR threshold was set at 5%. The complete proteomics dataset is available via ProteomeXchange (52, 53) with identifier PXD010576.

### Gene ontology analysis

Gene set enrichment analysis (GSEA) was run on the protein lists found to be significantly enriched at least 1.5 times in T0-T15, T0-T30 and T15-T30 using the online molecular signature database (MSigDB (23)) v6.2. Significantly enriched gene ontology (GO) pathways related to relevant “biological process” were extracted with their false-discovery rate (FDR). The Table S1 summarizes the relevant GO pathways associated to the T0-T15 time points. No significant enrichment was found at T0-T30 and T15-T30.

### Fluorescence microscopy

50 000 cells were plated on 24 well plates containing 12 mm cover glasses (Electron Microscopy Sciences) and incubated 48 h in complete medium containing 1 μg /ml of avidin. For the different eRUSH assays, cells were incubated at 5 min, 7 min, 12 min, 15 min and 30 min in complete medium containing 40 μM of biotin. Cells were fixed with 4% PFA for 20 min at room temperature and were permeabilized for 15 min with PBS containing 0.1% TritonX100 (Sigma-Aldrich), 0.5% bovine serum albumin (Euromedex). Cells were subsequently incubated one hour at room temperature with different primary antibodies (antibodies section), then one hour with secondary antibodies and DAPI staining. Cells were mounted with mowiol 4-88 (Sigma Aldrich). For LC3 staining, cells were fixed with formalin (Sigma-Aldrich) for 15 min at RT then with cold methanol for 5 min at −20°C, prior antibody staining in PBS containing 0.1% saponin and 1% FBS.

Images were taken with an AxioObserver Z1 inverted microscope (Zeiss) mounted with a CSU-X1 spinning disc head (Yokogawa), a back-illuminated EMCCD camera (Evolve, Photometrics) and a X63 (1.45 NA) or X100 (1.45 NA) oil objectives (Zeiss).

### Live imaging

About 250,000 cells seeded on 35 mm #1.5 glass bottom dishes (Ibidi) or on 25 mm cover glasses (Electron Microscopy Sciences) were transfected using JetPrime (Polyplus Transfection) according to manufacturer’s instructions. The dish was placed on the microscope stage, maintained in a dark atmosphere-controlled chamber at 37°C and 5% CO_2_. Live cell imaging was performed using an AxioObserver Z1 inverted microscope (Zeiss) mounted with a CSU-X1 spinning disc head (Yokogawa), a back-illuminated EMCCD camera (Evolve, Photometrics) and a X100, 1.45 NA oil objective (Zeiss) controlled by VisiView v.3.3.0 software (Visitron Systems). For TIRF microscopy, live imaging was performed with a TIRF PALM STORM microscope from Nikon using a back-illuminated EMCCD camera (Evolve 512, Photometrics) and a X100 APO, 1.49NA oil objective controlled by Metamorph, and an iLas^2^ FRAP/TIRF module (BioVision Technologies). The TIRF angle was chosen to obtain a calculated evanescent field depth < 100 nm.

### Preparation of protein extracts

Cells were seeded at 1.5 10^6^ cells per 10 cm dish per conditions in complete medium containing 1 µg/ml of avidin for 48 h. After incubation with biotin for 0 min, 30 min or 24 h, cells were washed 3 times with ice cold PBS and lysed with ice cold RIPA buffer (150 mM sodium chloride, 1% NP-40, 0.5% sodium deoxycholate, 0.1% sodium dodecyl sulfate (SDS), 50 mM Tris, pH 8.0, protease inhibitor (Promega). Cells were placed on ice for 10 min and span at 10,000 g for 20 min at 4°C. The supernatant was collected and subjected to the Pierce BCA assay kit (ThermoFisher Scientific).

### Western blot analysis

A total of 40 µg of protein lysates were run on Bolt 4-12% Bis-Tris plus gels (ThermoFisher Scientific) and proteins were transferred to nitrocellulose membranes. Nitrocellulose membranes were blocked with 5% (w/v) milk in PBS-T (PBS pH 7.4, 0.05% Tween 20) for 15 min. Primary antibodies (refer to antibody section) were incubated 1 h at RT or overnight at 4°C in PBS-T containing 5% milk. Secondary antibodies were incubated 1 h at room temperature. After washing with PBS-T, nitrocellulose membranes were incubated with Clarity Max western ECL substrate (Bio-Rad). The specific proteins were visualized with the ChemiDoc imaging system (Bio-Rad).

### Software analysis

Image processing was performed using either the FIJI upgrade of ImageJ (54) or the Imaris software v9.2 (Bitplane, Oxford Instruments). Quantifications for colocalization measurements were performed using Imaris software v9.2 (Bitplane, Oxford Instruments). Statistical analyses were performed with Microsoft Excel 2016 and Prism v7.04 (GraphPad). Flow cytometry analysis was done using the FlowJo software v10.4.2 (FlowJo, LLC). Raw mass spectrometry data were first analyzed using MaxQuant v 1.6.0.16. Differential proteomics data analysis was performed using DAPAR v1.10.3 and ProStaR v 1.10.4.

## Acknowledgements

We acknowledge the imaging facility MRI, member of the national infrastructure France-BioImaging. The mass spectrometry proteomics data have been deposited in the ProteomeXchange Consortium database (52, 53) with the identifier PXD010576. We thank Dr. Lucille Espert and colleagues for helpful discussions and sharing reagents related to autophagy. We thank Pr. Tom Kirchhausen for supportive discussions. This work was financially supported by the “Agence Nationale de la Recherche” (ANR) and the French Proteomic Infrastructure (ProFI; ANR-10-INBS-08-03). This work has been published within the framework of IdEx Université de Strasbourg and has received funding from the French State via the French National Research Agency (ANR) as part of the program “Investissements d’avenir” to R.G. This work was supported by an ATIP-AVENIR starting grant to R.G.

## Author contribution

MD and RG conceived the experiments. MD, IC, CD, VL and RG generated and characterized the TfR-eRUSH cell line. IC and RG performed flow cytometry. IC conducted the siRNA-based assays. MD and RG performed the microscopy analyses. FD and AH conducted mass spectrometry an FD, SC, CC and RG analyzed the proteomics data. ES and TX generated constructs for imaging and APEX labeling. GB and FP provided technical and conceptual support. RG and MD wrote the manuscript. RG, MD, CD, FD and CC edited and commented on the manuscript.

## Supporting information

**Fig. S1. Characterization and distribution of TfR-eRUSH over time post-biotin addition.**

(A-B) The western blot represents TfR expression in edited cells (TfR-eRUSH) compared to non-edited cells (wild type). Two molecular weight markers were used LMW (low molecular weight) and HMW (high molecular weight) to confirm TfR molecular size. Actin was used as a loading control. Anti-TfR was used in (A) and Anti-GFP in (B). (C) TfR-eRUSH cells were incubated for indicated times with biotin, fixed and stained with anti-calnexin antibodies (ER marker), anti-GM130 (cis-Golgi marker), anti-TGN46 (TGN marker) and revealed with appropriate secondary antibodies. The snapshots of cropped merged images from a single plane show TfR-eRUSH (green), Dapi staining (blue) and the indicated organelle marker (magenta). (D) Quantification of TfR-eRUSH colocalization with calnexin, GM130 or TGN46 at indicated times post-biotin addition. The Pearson’s correlation coefficient in the total volume of the cell was measured using the Imaris software. The graph shows mean Pearson’s correlation coefficient +/- SD and n = 17-20 cells from 2 independent experiments. Statistics were measured using unpaired t-test. * p value < 0.05, ** p value < 0.01, *** p value < 0.001. ns = non-significant.

**Fig. S2. TfR-eRUSH co-distributes with TMED10 but does not transit through mitochondria.**

(A) Representative images from a single z-stack indicate the localization of TfR-eRUSH relative to the endogenous TMED10 protein. TfR-eRUSH cells were treated for 12 min with biotin and images were acquired with a spinning disk confocal microscope. Representative images from a single plane show TfR-eRUSH (green), TMED10 (magenta) and nucleus (blue). Scale bar = 10 µm. Zoomed regions from white squares indicate a vesicle with TfR-eRUSH co-distributing with TMED10. Scale bar = 1 µm. (B) TfR-eRUSH cells were incubated for indicated times with biotin and mitochondria were visualized with 100 nM Mitotracker. Representative images from a single plane show TfR-eRUSH (green), mitochondria (magenta) and nucleus (blue). White dashed squares represent zoomed regions (lower panel). (C) Live imaging of TfR-eRUSH cells incubated with MitoTracker and biotin was performed at 30 s per frame using a spinning disk confocal microscope. An isolated event of TfR-mitochondria colocalization was observed 6 min post-biotin addition, although our spatial resolution limits the extend of this observation. The red square represented zoomed regions (left panel).

**Fig. S3. Rab7A vesicles transporting TfR-eRUSH does not contain Rab6A.**

(A) TfR-eRUSH cells transfected with Ruby3-Rab6A were imaged by TIRF microscopy 24 h post-transfection. Representative images were extracted at 10, 12 and 13 min post-biotin addition (scale bar = 10 µm). The white arrows show the Ruby3-Rab6A vesicles co-distributing with TfR-eRUSH. The yellow arrows indicate the zoomed regions of a Ruby3-Rab6A vesicle (magenta) co-distributing with TfR-eRUSH (green). Scale bar = 1 µm. (B) TfR-eRUSH cells were transfected with Ruby3-Rab7A for 24 h. Biotin was added for 15 min and images were taken with the spinning disk confocal microscope. Two cells were represented as images from a single z-stack. The staining indicates TfR-eRUSH (green), Ruby3-Rab7A (magenta), endogenous Rab6 (cyan), and the merge. The white arrows show Rab7A vesicles containing TfR-eRUSH but no Rab6. Scale bars = 5 µm.

**Fig. S4. TfR-eRUSH is not degraded following biotin addition.**

(A-B) Western blots representing the absence of degradation of TfR-eRUSH in presence of cycloheximide. Wild type and TfR-eRUSH cells were treated for 4 h with or without 50 µg/ml of cycloheximide and subsequently treated for 30 min with biotin. Actin was used as a loading control. Anti-TfR antibodies were used for visualization and HMW protein marker was used for molecular weight estimation in (A). Anti-GFP antibodies were used for visualization and LMW protein marker was used for molecular weight estimation in (B).

**Fig. S5. siRNA screen carried out on TfR-eRUSH cells**

Effect of Rab silencing on TfR-eRUSH expression. TfR-eRUSH cells were treated with siRNA sequences targeting 12 different Rab mRNAs and a non-targeting siRNA control. After 48 h post-transfection, the amount of total TfR-eRUSH at time 0 min was monitored by flow cytometry using EGFP fluorescence intensity. Anova test was run to assess for significance (* p value < 0.05, ** p value < 0.01, *** p value < 0.001).

**Table S1. Pathway enrichment analysis of proteins binding to TfR-eRUSH membranes at T0-T15.**

LC-MS/MS identification of the proteins associated to TfR-eRUSH membranes enriched > 1.5-fold at T15 compared to T0 and their associated ontology pathways.

**Table S2. Analysis of the Rab family members identified by mass spectrometry.**

LC-MS/MS identification of the proteins from the Rab GTPase family associated to TfR-eRUSH membranes. Fold change is shown for all detected Rabs. The red Rabs indicate that they are significantly enriched at T15 compared to T0.

**Table S3. List of the target sequences used in the siRNA screen.**

Oligonucleotides contained in the custom-made siGenome Smart pool cherry-pick library and used for the siRNA screen.

**Movie S1. Transport of neosynthesized TfR toward the PM.**

TfR-eRUSH cells were incubated with biotin and imaged using 3D spinning disk confocal microscopy. The images correspond to a z-stack spaced by 0.6 µm acquired every 30 s for 40 min. Three-dimensional reconstruction was performed using Imaris.

**Movie S2. TfR-eRUSH trafficking is mostly independent of the mitochondrial distribution.**

TfR-eRUSH cells were incubated with biotin and MitoTracker then imaged using 3D spinning disk confocal microscopy. The movie corresponds to a single plane from a z-stack spaced by 0.5 µm acquired every 10 s for 10 min. TfR-eRUSH is shown in green and MitoTracker in magenta. Scale bar = 2 µm.

**Movie S3. A subset of neosynthesized TfR-eRUSH traffics through Rab7-positive vesicles.**

TfR-eRUSH cells were transfected with Ruby3-Rab7A for 24 h and imaged in the presence of biotin using 3D spinning disk confocal microscopy. The movie corresponds to a single plane from a z-stack spaced by 0.3 µm acquired every 10 s from 7 min to 24 min. TfR-eRUSH is shown in green and Ruby3-Rab7A in magenta. Scale bar = 2 µm.

**Movie S4. TfR-eRUSH vesicles transiently interact with Rab7A.**

TfR-eRUSH cells were transfected with Ruby3-Rab7A for 24 h and imaged in the presence of biotin using TIRF microscopy. The movie represents a cropped vesicle starting from 706 s to 748 s post-biotin addition. A single evanescent field is acquired in TIRF mode every 2 s. TfR-eRUSH is shown in green and Ruby3-Rab7A in magenta. Scale bar = 2 µm.

**Movie S5. TfR-eRUSH vesicles interact with Rab6A.**

TfR-eRUSH cells were transfected with Ruby3-Rab6A for 24 h and imaged in the presence of biotin using TIRF microscopy. The movie represents a cropped vesicle starting from 838 s to 878 s post-biotin addition. A single evanescent field is acquired in TIRF mode every 2 s. TfR-eRUSH is shown in green and Ruby3-Rab6A in magenta. Scale bar = 2 µm.

