## Supplementary figures and images for "CRISPR-based bioengineering of the Transferrin Receptor revealed a role for Rab7 in the biosynthetic secretory pathway"

### Suppl Figure S1-S5

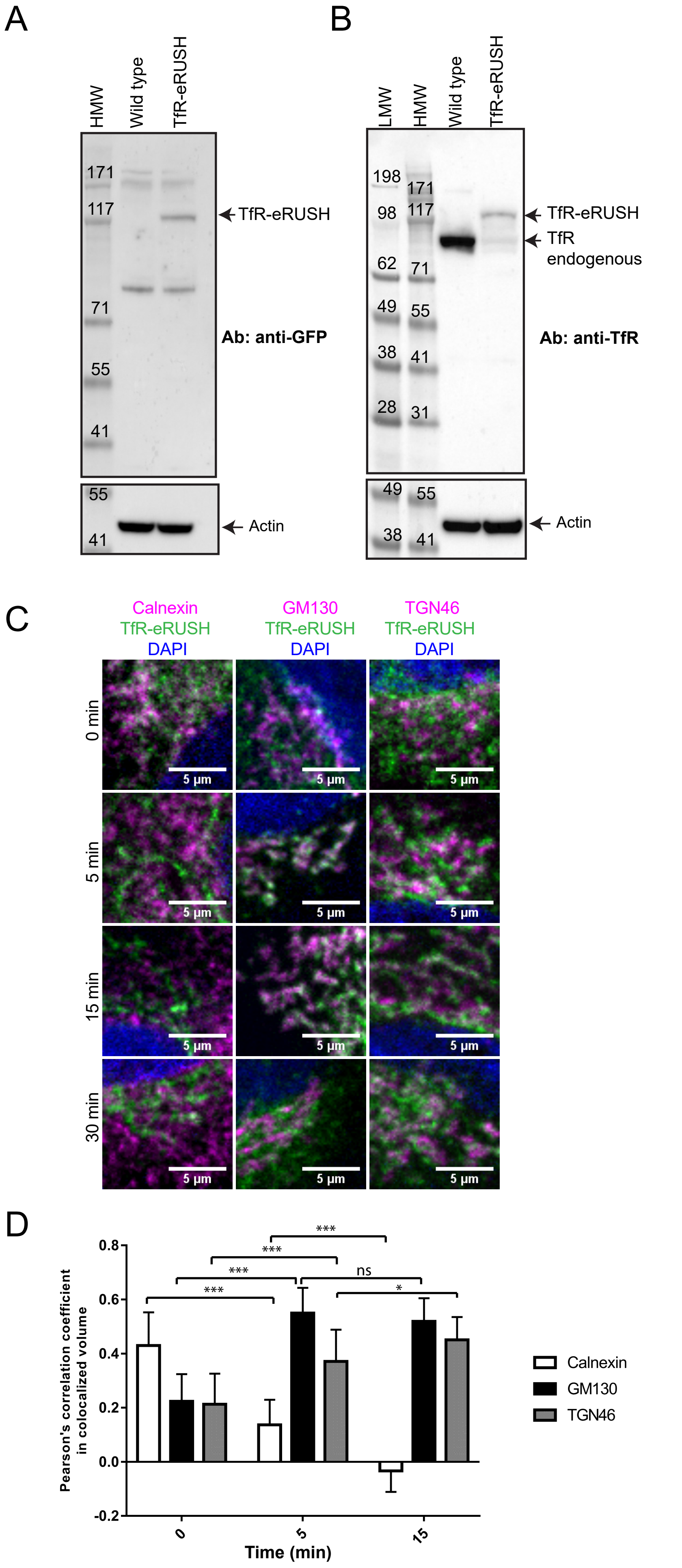


Fig S1. Deffieu et al.

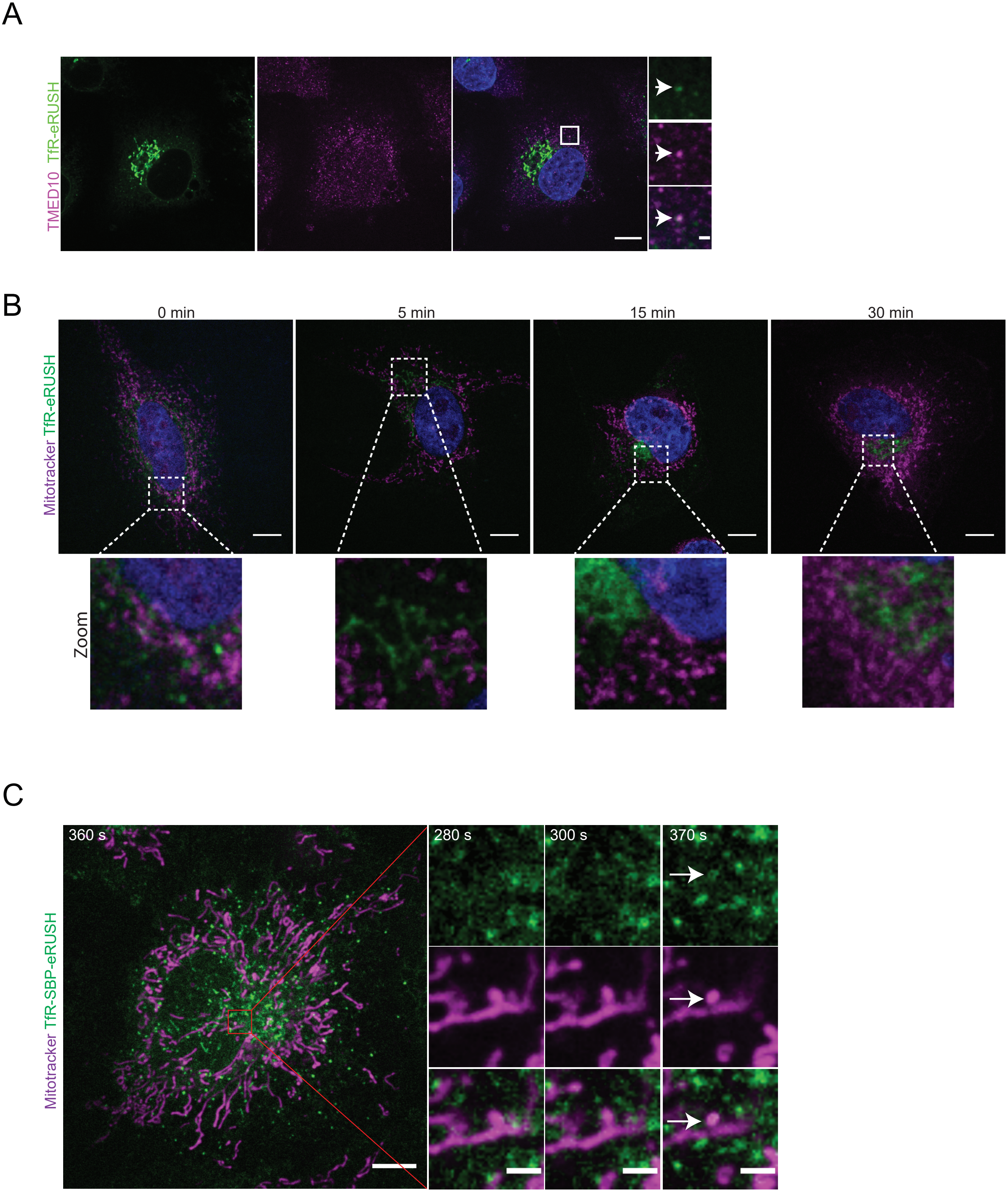


Fig S2. Deffieu et al.

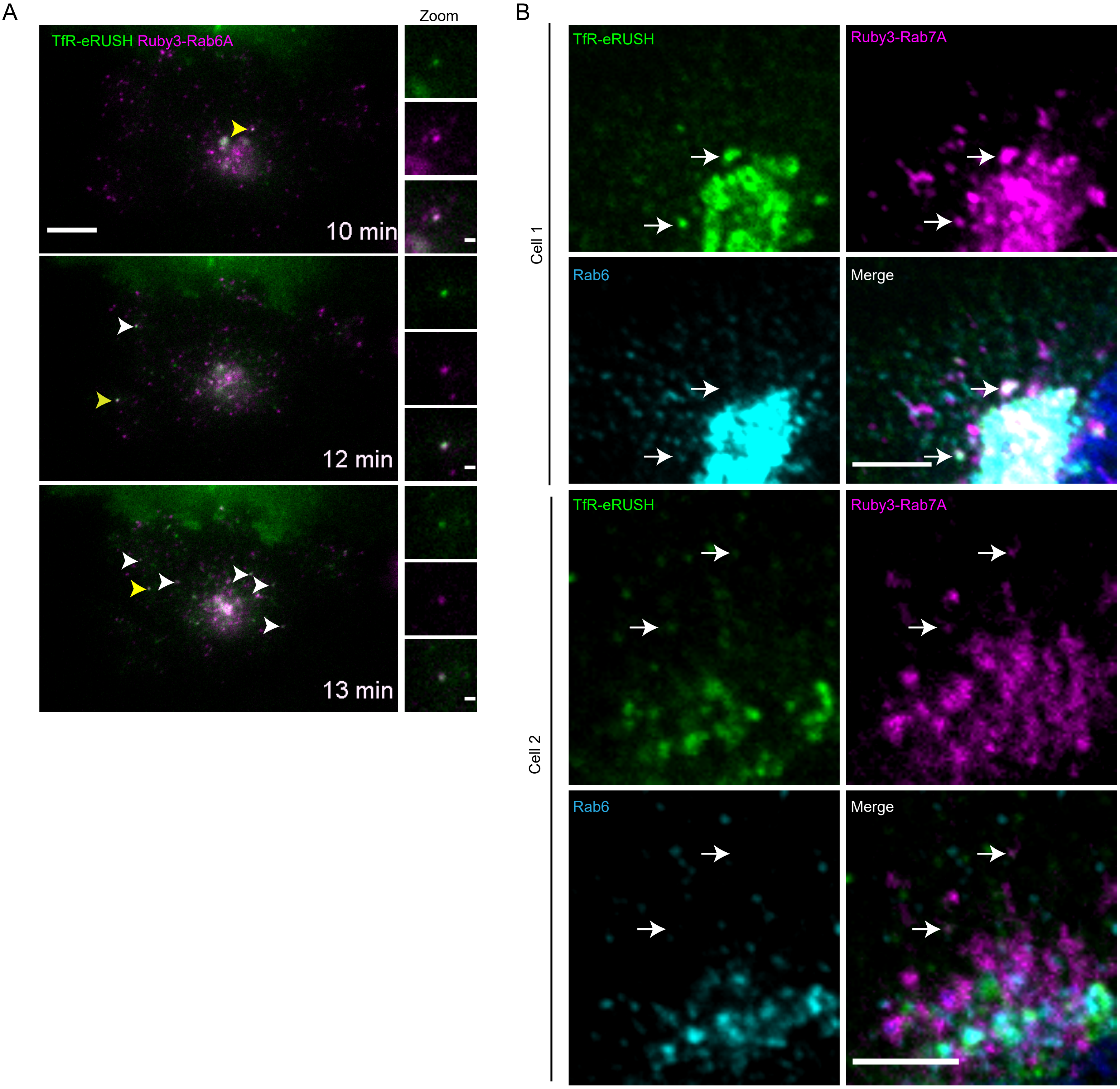


Fig S3. Deffieu et al.

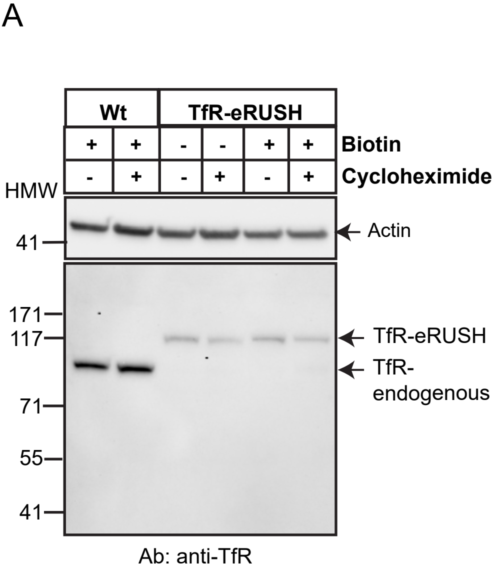


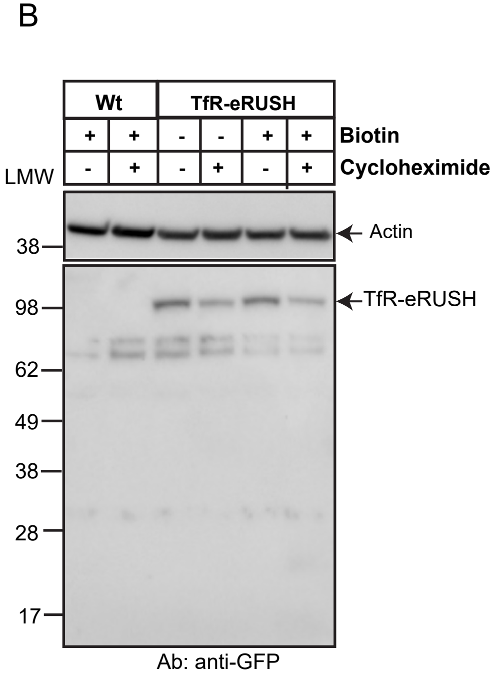


Fig S4 Deffieu et al.

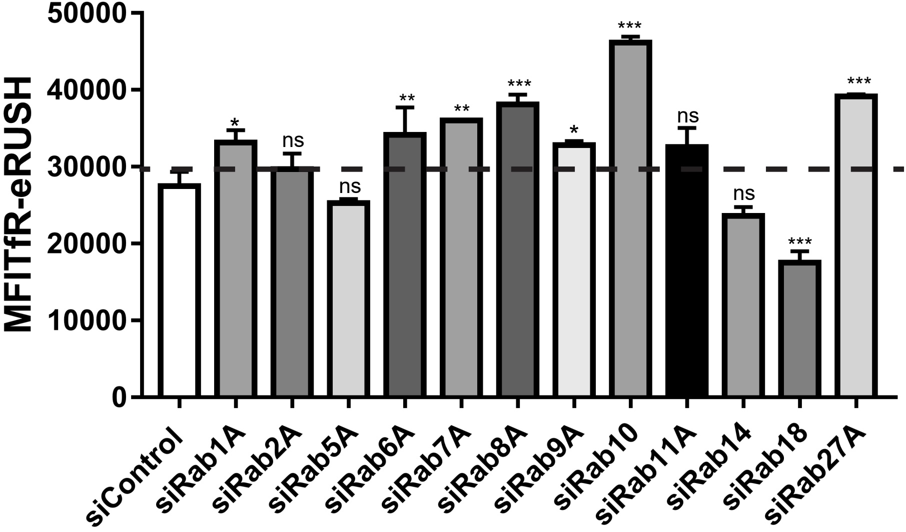


Fig S5. Deffieu et al.
